# Diagnostic, Infection Timing and Incidence Surveillance Applications of High Dynamic Range Chemiluminescent HIV Immuno-Assay Platforms

**DOI:** 10.1101/132332

**Authors:** Eduard Grebe, Alex Welte, Jake Hall, Sheila M. Keating, Shelley N. Facente, Kara Marson, Jeffrey N. Martin, Susan J. Little, Matthew A. Price, Esper G. Kallas, Michael P. Busch, Christopher D. Pilcher, Gary Murphy, on behalf of the Consortium for the Evaluation and Performance of HIV Incidence Assays (CEPHIA)

## Abstract

**Background:** Custom staging assays, including the Sedia HIV-1 Limiting Antigen Avidity EIA (LAg) and avidity modifications of the Ortho VITROS anti-HIV-1+2 and Abbott ARCHITECT HIV Ag/Ab Combo assays, are used to identify ‘recent’ infections in clinical settings and for cross-sectional HIV incidence estimation. However, the high dynamic range of chemiluminescent platforms allows differentiating recent and longstanding infection on signal intensity, and this raises the prospect of using unmodified diagnostic assays for infection timing and surveillance applications.

**Methods:** We tested a panel of 2,500 well-characterised specimens with estimable duration of HIV infection with the three assays and the unmodified ARCHITECT. Regression models were used to estimate mean durations of recent infection (MDRI), context-specific false-recent rates (FRR) and correlation between signal intensity and LAg measurements. A hypothetical epidemiological scenario was constructed to evaluate utility in surveillance applications.

**Results:** Over a range of MDRIs (reflecting recency discrimination thresholds), a diluted ARCHITECT-based RITA produced lower FRRs than the VITROS platform (FRR ≈ 0.5% and 1.5% respectively at MDRI of 200 days) and the unmodified diagnostic ARCHITECT produces incidence estimates with comparable precision to LAg (RSE ≈ 17.5% and 15% respectively at MDRI of 200 days). ARCHITECT S/CO measurements were highly correlated with LAg ODn measurements (r = 0.80) and values below 200 are strongly predictive of LAg recency and duration of infection less than one year.

**Conclusions:** Low quantitative measurements from the unmodified ARCHITECT obviate the need for additional recency testing and its use is feasible in clinical staging and incidence surveillance applications.

## Introduction

Laboratory assays for the detection, staging and management of HIV have been a significant priority over the course of the HIV/AIDS epidemic, with a great deal of innovation. In particular, the idea of using a test for ‘recent’ HIV infection to generate incidence estimates from cross-sectional surveys has attracted substantial attention and investment (1–11). A variety of candidate immunological and virological markers have been investigated, with most early applications focused on the expansion of the dynamic range of serological assays used to identify recent infection (*i.e.* increasing the length of time an incident case could still be classified as recently infected). In conventional plate reader-based ELISA platforms, high diagnostic sensitivity requires a rapid rise of signal strength with increasing antibody titre and avidity, so that patients’ readings rapidly traverse the range of quantifiable detection as the infection progresses. Customisations to expand the dynamic range and facilitate the introduction of a reproducible recent/non-recent threshold have variously taken the form of dilution, incubation time reduction, or antibody-antigen binding degradation (sometimes in combination) to produce markers indicative of antibody titre, HIV-specific proportion, or avidity.

Individuals who are virally suppressed (either through effective endogenous control or by antiretroviral treatment) tend to undergo at least partial seroreversion, leading to ‘false’ recent classifications under protocols designed for cross-sectional incidence estimation. Inspired by the hypothesis that antibody *quality* reverts less than antibody *titre* or *proportion*, the notion of a ‘two-well avidity modification’ has gained some traction as a promising generic approach. In this case a specimen is subjected to two runs on the diagnostic platform, under different conditions: an ‘untreated’ (*i.e.* close to standard conditions) reaction well and a ‘treated’ reaction well, in which conditions are altered to restrict antibody-antigen binding. The ratio of the signals generated by ‘treated’ *vs.* ‘untreated’ reaction wells – usually named an ‘avidity index’ – is interpreted as a measure of antibody binding capacity. Avidity typically increases over time after infectious exposure as the antibody response matures.

Two-well avidity modifications of two modern chemiluminescent platforms with high intrinsic dynamic range, the Ortho VITROS anti-HIV-1+2 Assay (12) and the Abbott ARCHITECT HIV Ag/Ab Combo Assay (13), have been proposed. These employ ‘chaotropic’ agents – *i.e.* agents that interfere with antibody-antigen binding – in the ‘treated’ reaction well to inhibit the formation of antibody-antigen complexes or disrupt complexes after formation. While not relying on specimen dilution (which is an alternative, non-avidity approach to dynamic range expansion), the use of an additional reagent in the modified run requires some dilution of samples in the ‘untreated’ run to match input volumes and render the resulting signals comparable.

Performance of the VITROS and ARCHITECT avidity assays for use in cross-sectional incidence estimation has been investigated by the CEPHIA collaboration, as part of an independent evaluation of leading candidates for HIV incidence assays (9,10). During application of the proposed procedures for recency ascertainment using the CEPHIA ‘evaluation panel’, these two-well avidity protocols additionally produced results from diluted (1:10) but otherwise unmodified runs. These previously unreported data, and a new evaluation of the unmodified Abbott ARCHITECT HIV Ag/Ab Combo Assay (*i.e*., applied according to the manufacturer’s Instructions for Use [IFU] in diagnostic applications) by CEPHIA form the subject of this work. We further report, for comparison, results from the Sedia™ HIV-1 Limiting Antigen Avidity (LAg) assay on the same CEPHIA evaluation panel.

Until recently, no practical recent/non-recent infection classification scheme has been constructed using only markers available through unmodified diagnostic platforms and algorithms. This is a noteworthy opportunity, given that the Western blot, at times widely used as a ‘confirmatory’ diagnostic test, provides a compelling picture of immune response evolution (14). In many settings, it has been noted that ‘indeterminate’ Western blot patterns tend to evolve into unambiguously HIV-positive patterns within a few weeks. However, lack of consistent production and quantitation of Western blot band intensity has been one key limitation preventing use of the Western blot as a staging assay. Similarly, the cost and complexity of using customised recency assays in disease surveillance has limited adoption (15), and the potential to interpret data already routinely available from diagnostic assays to classify infections as recent/non-recent may improve the feasibility of incidence surveillance in many settings.

In clinical settings, the only well-known early infection staging scheme is ‘Fiebig staging’ (14), nominally based on unmodified commercial assays, but using only the qualitative/categorical results. ‘Fiebig stages’ are based on discordance between tests of varying diagnostic sensitivity, or, more precisely, the mean time from infectious exposure to test conversion. Given progress in the reduction of diagnostic delays (window periods), such a staging system has utility only in the very early stages of infection, and inferences about the duration of infection can in practice only be made when specific diagnostic platforms, many of which are no longer available, are employed.

We report the first evidence that certain unmodified chemiluminescent 4th generation diagnostic assays can provide meaningful recent/non-recent infection staging information applicable to incidence surveillance, or for clinical interpretation in post-diagnosis counselling. The latter application is already being practised in several countries where custom recency staging assays (such as LAg) are applied to specimens from newly diagnosed persons. This involves additional laboratory resources and longer turnaround times, as specimens are usually reflexed to a small number of centralised specialist laboratories for recency ascertainment.

## Methods

### The CEPHIA Specimen Repository and Evaluation Panel

The CEPHIA specimen repository houses over 25,000 HIV-1-positive specimens. Specimen details and background clinical data (obtained from contributing clinical cohort studies) are stored in harmonised form in a research database. As previously described (9,10,16), the CEPHIA Evaluation Panel consists of 2,500 plasma specimens, selected to allow a full independent assessment of promising tests for recent infection, including estimation of test properties relevant to HIV incidence estimation. The specimens were obtained from 932 unique subjects (1-13 specimens per subject). Most specimens were obtained from subjects infected with subtype B (53% of specimens), subtype C infection (27%), subtype A1 (12%) and D (6%). The panel further contained multiple blinded aliquots of three control specimens (25 replicates of each) with antibody reactivity characteristic of recent, intermediate and longstanding infection to allow evaluation of the reproducibility of assay results. The majority (67%) of subjects contributing specimens to the panel had sufficient clinical data to produce Estimated Dates of Detectable Infection (EDDIs), which are obtained by systematically interpreting diverse diagnostic testing histories by means of the ‘diagnostic delays’ – the average time elapsed from exposure to HIV to a first positive result on the assay in question (11).

The UCSF Human Research Protection Program & IRB (formerly CHR, #10-02365) approved study procedures.

### Laboratory procedures

For the present analysis, data for the VITROS Avidity, ARCHITECT Avidity and Sedia LAg Avidity assays were generated as previously described (9,10), with the exception that the data from the CEPHIA evaluations of the VITROS Avidity and the ARCHITECT Avidity assays were reanalysed using only the ‘untreated’ (diluted) run (*i.e.* the sample diluted 1:10 with PBS). Additional testing of the CEPHIA evaluation panel was undertaken according to the manufacturer’s IFU for the Abbott ARCHITECT HIV Ag/Ab Combo assay. Data for the LAg assay is included for comparison.

Specimens were independently tested in CEPHIA laboratories (Blood Systems Research Institute, San Francisco, CA – VITROS Avidity and National Infection Service, Public Health England, London, UK – ARCHITECT Avidity, unmodified ARCHITECT, Sedia LAg) by technicians blinded to specimen background data.

The procedures used in the testing of the two avidity modifications and LAg have been previously described (9). The unmodified ARCHITECT HIV Ag/Ab Combo assay is a two-step immunoassay which detects both the presence of HIV p24 antigen and antibodies to HIV-1 and HIV-2 in human serum and plasma. For this evaluation, we only investigated anti-HIV-1 detection. In the first step, the sample, assay diluent, and paramagnetic microparticles are combined. HIV p24 antigen and HIV-1/HIV-2 antibodies present in the sample bind to the HIV-1/HIV-2 antigen and HIV p24 monoclonal (mouse) antibody-coated microparticles. After washing, the HIV p24 antigen and HIV-1/HIV-2 antibodies bind to the acridinium-labelled conjugates (HIV-1/HIV-2 antigens, synthetic peptides, and HIV p24 antibody). Following another wash cycle, pre-trigger and trigger solutions are added to the reaction mixture. The resulting chemiluminescent reaction is measured as relative light units (RLUs). The assay incorporates a number of controls and calibrator specimens, and regular maintenance of the ARCHITECT platform is required to ensure accuracy of results. For the purpose of this evaluation specimens were tested and analysed in singleton but for diagnostic purposes the manufacturer suggests specimens be tested in duplicate.

### Statistical Analysis

For use in incidence surveillance, the performance of recent infection tests is summarised by two parameters, the Mean Duration of Recent Infection (MDRI) and the False-Recent Rate (FRR).

*MDRI* denotes the average amount of time that individuals spend exhibiting the ‘recent’ biomarker, while infected for less than some cut-off time (denoted *T*, 2 years in the present work). This captures the defining biological aspects of the recency test, and should be more than approximately half a year to yield informative incidence estimates from feasibly-sized surveys, even in high incidence settings. As described previously (3, 4), MDRI was estimated by fitting a linear binomial regression model for the probability of testing recent as a function of estimated time since infection, *P*_*R*_(*t*), using a logit link function and a cubic polynomial in time (since estimated date of detectable infection). The function was fit to data points up to 800 days post-infection. The MDRI was then obtained by integrating the function from 0 to *T*. Confidence intervals were obtained by resampling subjects in 10,000 bootstrap iterations.^1^

The *FRR* is the proportion of those individuals who are infected for longer than the cut-off time *T*, but who nevertheless produce a recent result on the test. For surveillance, values of FRR above approximately 1-2% are highly vulnerable to bias and artefacts during the incidence estimation procedure.

FRR is inevitably context-dependent. In order to estimate FRR, a hypothetical epidemiological scenario was constructed and the FRR in untreated and treated individuals was estimated separately and weighted according to the treatment coverage specified in the scenario. The epidemiological scenario used in the present study can be summarised as: 1) HIV prevalence of 30%; 2) incidence of 1.5 cases per 100 person-years; and 3) treatment coverage (defined as proportion of HIV-infected individuals who are on ART, all of whom are assumed to be virally suppressed) of 80%.

To estimate the FRR in untreated individuals, the function *P*_*R*_(*t*) was fit using data from all times post-infection, and weighted according to the probability density function for times since infection in the untreated population. The distribution of times since infection was parameterised as a Weibull survival function, with the shape and scale parameters chosen to produce the desired treatment coverage in a population with the specified incidence and prevalence, and normalised to the specified recent incidence. The FRR in treated subjects, *P*(*R*|*tx*) is simply the binomially estimated probability that treated subjects infected for longer than *T* would produce a recent result.^2^

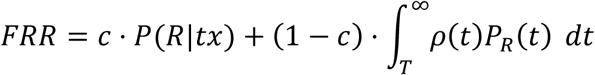

where *c* is the treatment coverage, 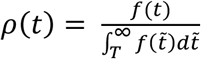 and 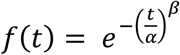

For surveillance applications, utility is defined in terms of the standard error on the incidence estimator (17). To assess this, MDRI and context-specific FRR were calculated for a range of ‘recent/non-recent’ discrimination thresholds, and the variance on the incidence estimator was calculated for the specified epidemiological context (assuming a demonstrative simple random sample of 10,000 individuals). A key difference between a previously constructed scenario (10) and the present analysis is in the HIV-positive case definition, which previously required a fully-developed Western blot, and in the present case is expanded to all individuals who test positive on a qualitative NAT assay (threshold of detection 30 HIV-RNA copies/mL). This has ramifications for MDRI, which was estimated using EDDIs in which detectable infection is defined as positivity on a hypothetical viral load assay with a detection threshold of 1 copy/mL. Reported MDRI estimates must be interpreted against this reference standard, and for the purposes of incidence estimation, MDRI was adjusted to account for the sensitivity of the HIV screening algorithm (*i.e*., 4.8 days shorter than reported MDRI to account for the diagnostic delay of the Aptima HIV-1 RNA Qualitative Assay used in NAT screening).

MDRI, FRR and the variance of the incidence estimate in the hypothetical scenario were computed using the R package *inctools* (18).

Linear regression was used to assess the correlation between ARCHITECT S/CO readings and LAg ODn readings on the untreated subset of the evaluation panel. Binomial logistic regression models were employed to assess the predictive value of ARCHITECT S/CO values for (1) duration of infection ≤ one year, and (2) LAg ODn values ≤ 1.5 (the conventional recency discrimination threshold). The logistic regression models had the following form:

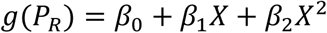

with *P*_*R*_ the probability of recency (duration of infection ≤ 1 year or LAg ≤ 1.5), *g*() the logit link function and *X* the ARCHITECT S/CO measurements.

## Results

Figure 1 shows, in four panels customised to the apparent dynamic range of each assay, the MDRI (y-axis) as a function of recent/non-recent discrimination threshold (x-axis, range chosen to yield MDRI values from 50 to 400 days). A viral load threshold of 100 HIV RNA copies/mL is applied in all cases. MDRI is tuneable by adjusting the recent/non-recent discrimination threshold, but FRR increases with increasing MDRI.

**Figure 1:**
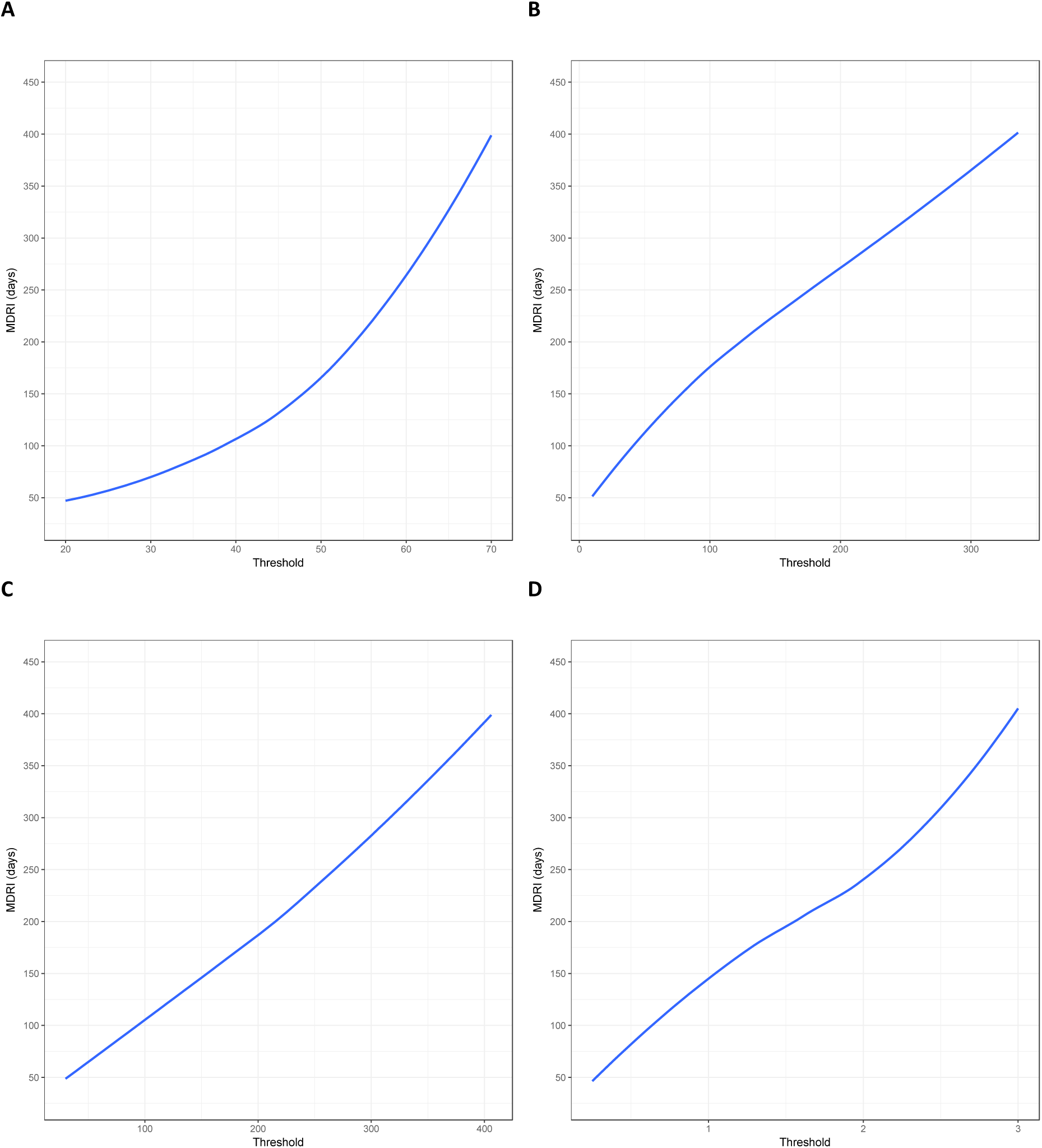
Mean Duration of Recent Infection by recency discrimination threshold. Mean Duration of Recent Infection (y axis), as a function of recency discrimination threshold (x axis, range chosen to yield MDRI values from approximately 50 to 400 days). A) Minimally diluted (‘untreated well’) of the Ortho VITROS anti-HIV-1+2 Assay avidity modification B) Minimally diluted (‘untreated well’) of the Abbott ARCHITECT HIV Ag/Ab Combo Assay avidity modification C) The completely unmodified (manufacturer IFU) Abbott ARCHITECT HIV Ag/Ab Combo Assay D) (As a reference) the Sedia HIV-1 Limiting Antigen Avidity EIA (LAg)

Low ARCHITECT S/CO measurements are highly ‘predictive’ of low LAg normalised optical density (ODn) measurements, as can be seen in Figure 2, which shows a scatterplot and linear regression model for LAg ODn vs. ACRHITECT S/CO (panel A) and binomial logistic regression for the probability of obtaining an ODn ≤ 1.5 (the conventional recent/non-recent discrimination threshold) and duration of infection less than one year (panel B). Table 1 further reports that 78% of untreated specimens in the evaluation panel with S/CO readings < 200 and viral load measurements > 100 copies/mL produce LAg ODn values ≤ 1.5, and 87% of these specimens were drawn within one year of EDDI. Low quantitative readings from the diagnostic assay, together with above-threshold viral load, therefore appear to obviate the need for additional staging assays.

**Table 1:**
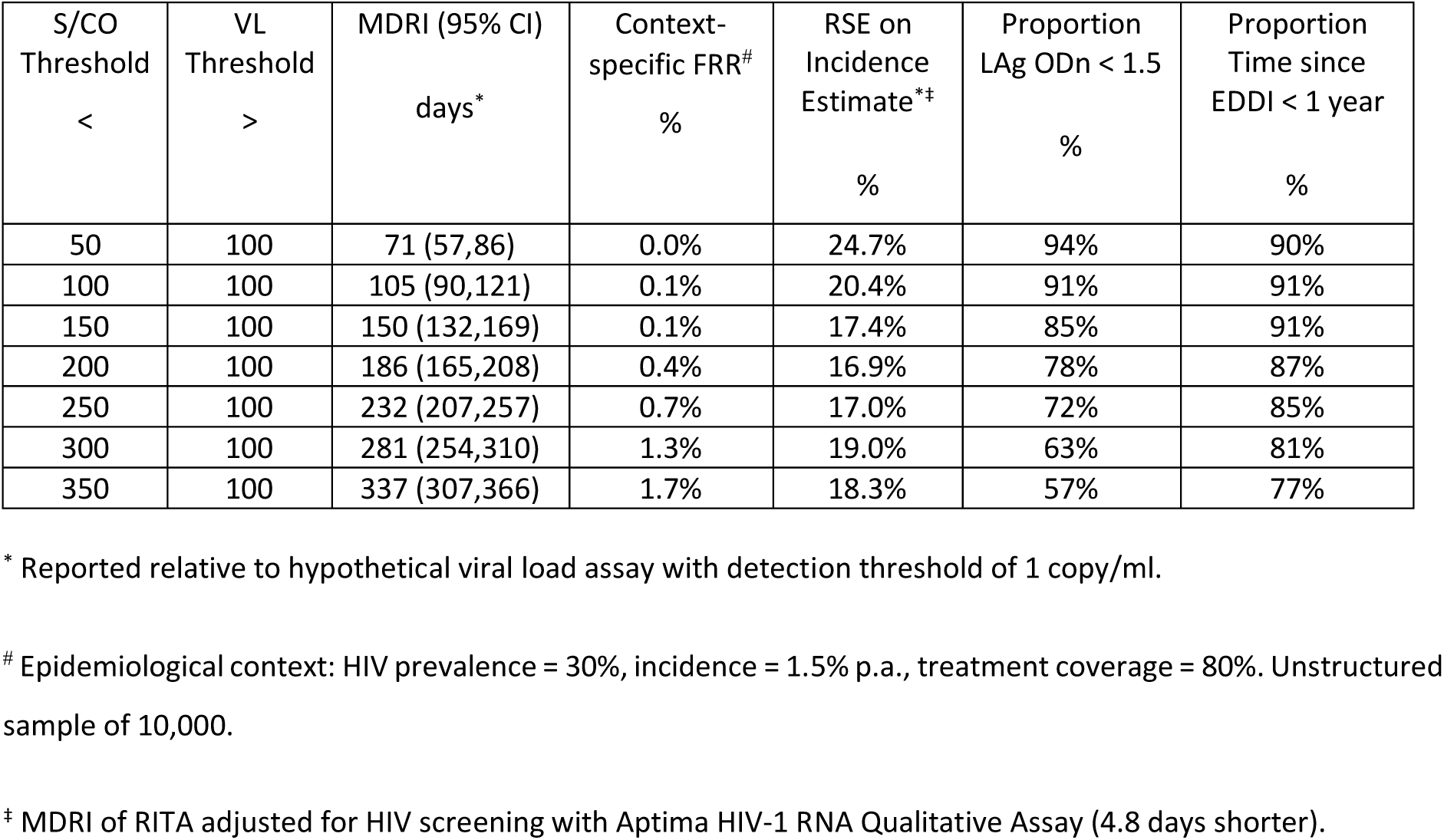
Performance characteristics of the unmodified ARCHITECT assay. Mean Duration of Recent Infection, context-specific False-Recent Rate (in the specified hypothetical epidemiological scenario), Relative Standard Error on the incidence estimate (in the specified hypothetical scenario^*#*^), proportion LAg-recent and proportion infected less than one year, by Signal-to-Cutoff Ratio recency discrimination threshold and viral load threshold.

| S/CO Threshold<br>< | VL Threshold<br>> | MDRI (95% CI)<br>days* | Context-specific FRR<sup>#</sup><br>% | RSE on Incidence Estimate*<sup>‡</sup><br>% | Proportion LAg ODn < 1.5<br>% | Proportion Time since EDDI < 1 year<br>% |
| --- | --- | --- | --- | --- | --- | --- |
| 50 | 100 | 71 (57,86) | 0.0% | 24.7% | 94% | 90% |
| 100 | 100 | 105 (90,121) | 0.1% | 20.4% | 91% | 91% |
| 150 | 100 | 150 (132,169) | 0.1% | 17.4% | 85% | 91% |
| 200 | 100 | 186 (165,208) | 0.4% | 16.9% | 78% | 87% |
| 250 | 100 | 232 (207,257) | 0.7% | 17.0% | 72% | 85% |
| 300 | 100 | 281 (254,310) | 1.3% | 19.0% | 63% | 81% |
| 350 | 100 | 337 (307,366) | 1.7% | 18.3% | 57% | 77% |

\* Reported relative to hypothetical viral load assay with detection threshold of 1 copy/ml.
<sup>#</sup> Epidemiological context: HIV prevalence = 30%, incidence = 1.5% p.a., treatment coverage = 80%. Unstructured sample of 10,000.
<sup>‡</sup> MDRI of RITA adjusted for HIV screening with Aptima HIV-1 RNA Qualitative Assay (4.8 days shorter).

**Figure 2:**
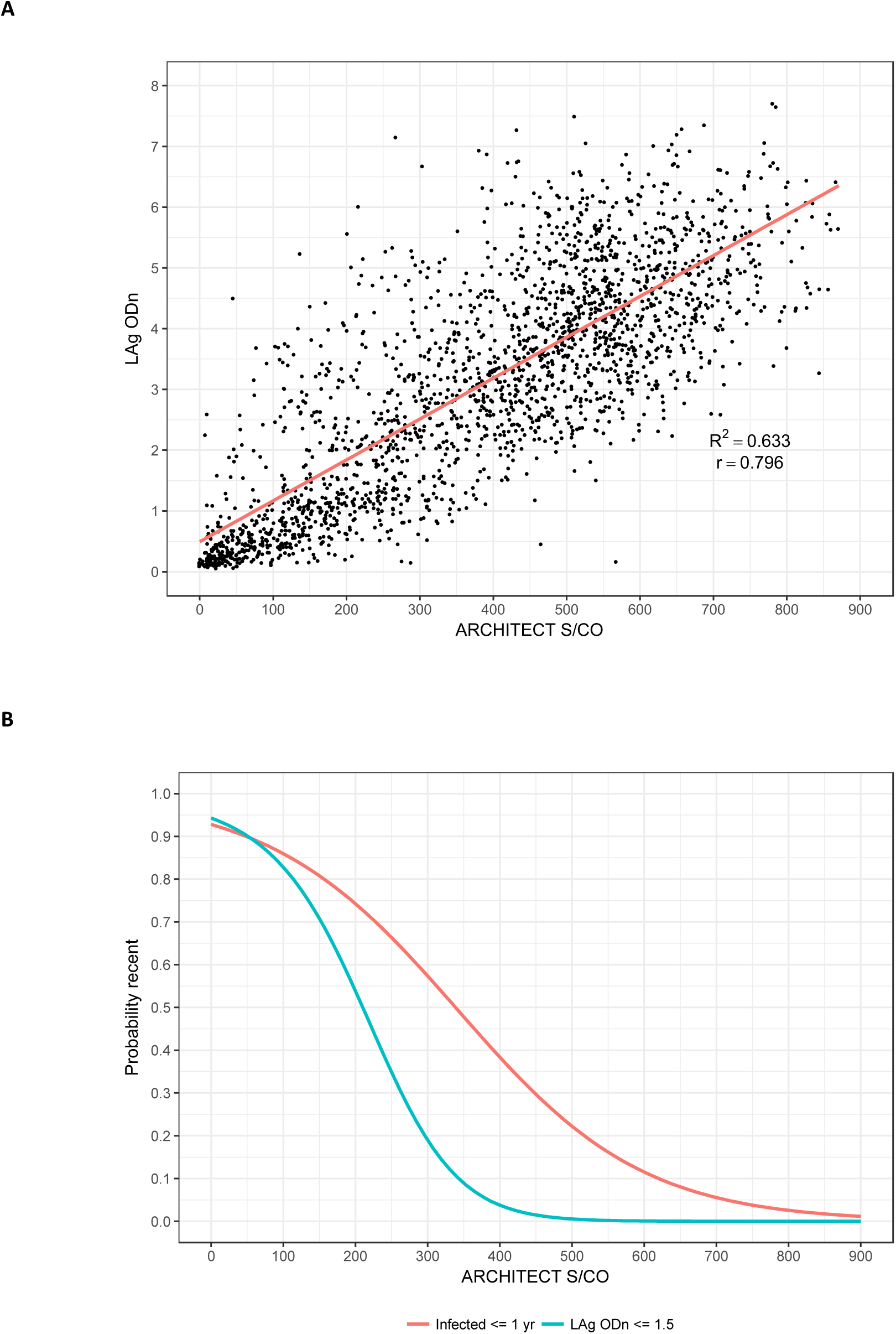
Low ARCHITECT S/CO measurements ‘predict’ low LAg ODn measurements and short duration of infection. A) Scatterplot of ARCHITECT (x-axis) and LAg (y-axis) readings, and linear regression model B) Binomial logistic regression models for the probability of obtaining a LAg ‘recent’ result (ODn ≤ 1.5) and probability of duration of infection ≤ 1 year

Figure 3 summarises performance metrics for the minimally diluted ‘untreated wells’ of the VITROS and ACRHITECT Avidity assays, the unmodified ARCHITECT diagnostic assay and LAg as part of a RITA in surveillance applications. Panel A shows context-specific FRR in the scenario outlined above against MDRI (encoding recency discrimination threshold) and panel B shows the relative standard error on the incidence estimate against MDRI. Comparison of the diluted VITROS and ACRHITECT platforms shows a much more rapid rise in FRR for the VITROS platform, reaching more than 5% when MDRI is approximately one year. For this reason, the ARCHITECT platform was selected for further evaluation in entirely unmodified form (manufacturer’s IFU). The results indicate that a RITA based on the unmodified ARCHITECT achieves an only marginally greater FRR than the diluted version, and that its performance for surveillance purposes (as captured in the precision of incidence estimate) is very similar to that of a RITA based on the widely-employed LAg assay.

**Figure 3:**
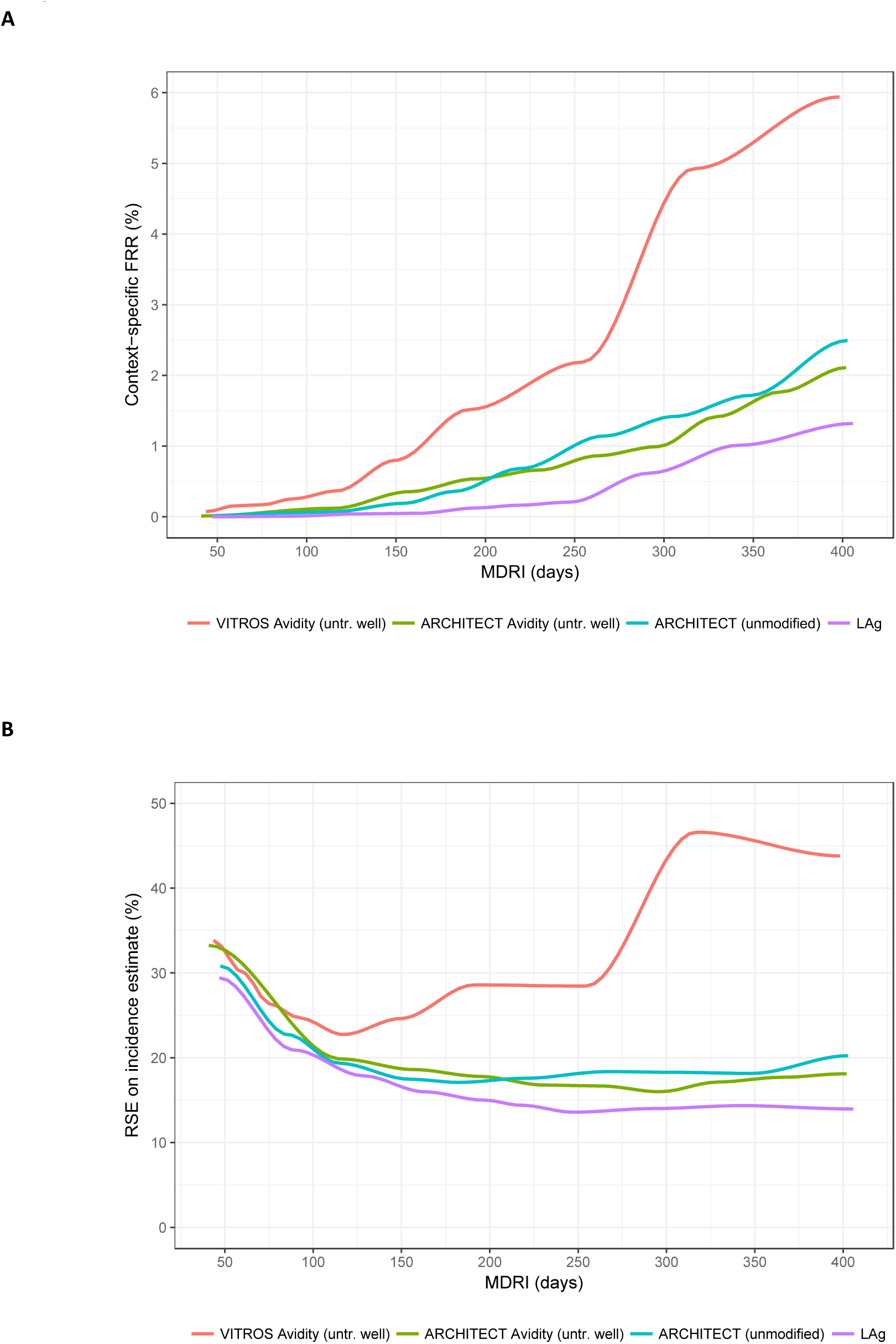
Performance characteristics by Mean Duration of Recent Infection. Context-specific False-Recent Rate (FRR) and relative standard error (RSE) on incidence estimate in a demonstrative epidemiological context (30% HIV prevalence, 1.5% per annum incidence, 80% treatment coverage) and using an unstructured sample of 10,000 respondents. A) Context-specific False-Recent Rate B) Precision of Incidence Estimate

Table 1 shows performance metrics for a range of S/CO thresholds: MDRI, context-specific FRR and relative standard error on an incidence estimate in the epidemiological context outlined above, with MDRI adjusted for the sensitivity of a practical screening algorithm. In this context, a S/CO threshold of 200 appears close to optimal, yielding an MDRI of 186 days (95% CI: 165,208), a context-specific FRR of 0.4%, and a relative standard error on the incidence estimate of 16.9%). S/CO thresholds < 100 or > 300 produce less precise incidence estimates in surveillance applications as a result of shorter MDRI and higher FRR respectively, but the lower threshold could be useful for clinical staging applications.

## Discussion

In Figure 1 we can see that, although the actual ranges of plausible thresholds vary considerably between platforms, the chemiluminescent diagnostic assays have sufficient dynamic range to support simple threshold-based definitions of recent infection. For surveillance applications, such as case-based surveillance where central laboratories apply comparable, or comparably well-calibrated, recency staging assays to a large number of cases identified in a health system; blood banking applications; and large population-based surveys. Optimal threshold choice depends on the specific epidemiological and methodological context of application, but would involve a similar trade-off between MDRI and FRR as that demonstrated in Table 1 and Figure 3. It is worth noting that even state-of-the-art staging algorithms exhibit disappointing performance in surveillance applications, with precise incidence estimates emerging only at extraordinarily high sample sizes or high values of incidence (10,19). Recency staging data is unlikely ever to be the only source of incidence-related information – meaning that it should be used in conjunction with estimation procedures (still under development) which also use the demographic (age and time) structure of prevalence data and mortality (20,21).

In interpreting data for individual use it is important to consider an inherent limitation of any diagnostic test: the potential for false positive results. While a diagnostic HIV interpretation would not be based on the outcome of a single test, the lack of confirmatory tests for ‘recent’ infection means that a single test is often used for recency determination. It is likely that a false positive or non-specific reaction would give a low S/CO reading, and thus would be more likely to be misclassified as a recent infection. Consideration must be given to confirming the antibody positive status of the individual. In addition, the very nature of combined antigen/antibody 4^th^ generation tests means that antigen-only positive samples may also be detected. These results, which are almost certainly truly recent infections, may also generate a low S/CO depending on a number of factors, including the amount of antigen present and whether this antigen has complexed with developing antibody, and should not be discarded over the absence of antibody reactivity. Given this, before interpreting a test for recency it is critical to ensure that the HIV diagnosis is confirmed by local testing algorithms and criteria that identify non-specific reactivity and differentiate antibody and antigen reactivity.

One must not dismiss the performance of the unmodified ARCHITECT demonstrated in the present analysis, simply because it is not quite as good as what can currently be achieved by other staging algorithms based on more complex protocols. Substantial logistical and cost advantages, in both routine contexts and population-based surveys, could be achieved by using a single serological assay for diagnostic and staging purposes, especially in the light of the high-throughput, automated nature of these platforms.

Estimating time since infection based on HIV biomarker progression is made difficult by the complex nonlinear growth exhibited by immune markers of disease progression, and the absence of a robust model for capturing inter-subject variability leads to an inability to formally measure the precision of the timing estimate. Nevertheless, simply ‘looking up’ the assay result on the threshold-MDRI curve, and noting that “*X* reactivity level of this diagnostic assay is one that subjects, on average, spend *Y* time beneath”, provides clinically meaningful information. In principle, the most natural and coherent way to provide realistic estimates for individual-level times since infection would involve the use of testing histories and other contextually-derived ‘prior information’ in a Bayesian framework. This would require a high level of confidence that the model of biomarker growth correctly captures complex inter-subject variability.

Critical clinical decisions – such as whether to offer antiretroviral treatment – could be based on the transformation, through a calibration curve of a test result into a timescale (only loosely interpreted as an estimate of time since infection). Typically, a simple “likely recent/non-recent” result is currently fed back to clinician and patient, qualified by a mean time meant by recent, with caveats outlining a number of known confounders of accurate assay performance. Our proposal uses the specific assay result, rather than simply dichotomising results into a recent/non-recent classification (based on a single threshold, which may be far from the particular patient’s result), and therefore offers an interpretation that is much more informative.

This analysis provides the first evidence that unmodified diagnostic assays can provide meaningful staging information applicable in clinical and surveillance settings. Even a recent analysis of the Bio-Rad Geenius^TM^ HIV1/2 supplemental assay (16), though using a platform available in diagnostic settings, requires a research-use-only modification of the cartridge reader software to facilitate the extraction of quantitative band intensities, which form the basis of the recency classification.

Tests that intrinsically serve both as part of a diagnostic algorithm and a staging algorithm offer obvious practical advantages, and perhaps more significantly, create a genuine market advantage. This may stimulate investment in the otherwise unattractive sector of staging assays (15). While customised staging assays will remain useful in many contexts, the ability to extract staging information from diagnostic assays constitutes a significant advance for the fields of clinical staging and incidence surveillance.

## Footnotes

1 Since the estimation procedure does not rely on longitudinal biomarker progression within individuals, the bootstrapping procedure resamples *subjects* rather than individual data points (with replacement) to account for the non-independence of measurements on specimens drawn from the same individual.

2 In the CEPHIA Evaluation Panel, all treated subjects were virally suppressed, resulting in an estimate of P(R|tx) = 0 in all cases where a supplemental viral load threshold is applied. In real-world populations, it is likely that a certain (unknown) proportion of treated subjects would be virally unsuppressed and that the FRR in treated subjects would therefore be non-zero.

